# The female urinary microbiota in relation to the reproductive tract microbiota

**DOI:** 10.1101/628974

**Authors:** Chen Chen, Lilan Hao, Weixia Wei, Fei Li, Liju Song, Xiaowei Zhang, Juanjuan Dai, Zhuye Jie, Jiandong Li, Xiaolei Song, Zirong Wang, Zhe Zhang, Liping Zeng, Hui Du, Huiru Tang, Huanming Yang, Jian Wang, Karsten Kristiansen, Xun Xu, Ruifang Wu, Huijue Jia

**Author notes:** These authors contributed equally to this work (Chen Chen, Lilan Hao, Weixia Wei). Correspondence should be addressed to H. J. or R. W. **E-mails and ORCIDs:** Chen Chen*., Lilan Hao*., Fei Li., Liju Song., Xiaowei Zhang., Zhuye Jie., Jiandong Li., Xiaolei Song., Zirong Wang., Zhe Zhang., Huanming Yang., Jian Wang, Karsten Kristiansen., Xun Xu, Huijue Jia†., Weixia Wei*., Juanjuan Dai., Liping Zeng., Hui Du., Huiru Tang., Ruifang Wu†.

## Abstract

Human urine is traditionally considered to be sterile, and whether the urine harbours distinct microbial communities has been a matter of debate. The potential link between female urine and reproductive tract microbial communities is currently not clear.

Here we collected the urine samples from 147 Chinese women of reproductive age, and explored the nature of colonization by 16S rRNA gene amplicon sequencing, real-time qPCR and live bacteria culture. To demonstrate utility intra-individual Spearman’s correlation was used to explore the relationship between urine and multi-sites of the reproductive tract. PERMANOVA was also performed to explore potential correlations between the lifestyle and various clinical factors and urinary bacterial communities. Our data demonstrated distinct bacterial communities in urine, indicative of a non-sterile environment. Types of diverse, *Streptococcus*-dominated, and *Lactobacillus*-dominated were the three most common types in the cohort. Detailed comparison of the urinary microbiota to the multi-sites of reproductive tract microbiota demonstrated the urinary microbiota was more similar to the microbiota in the cervix and uterine cavity instead of vagina in the same women.

Our data demonstrates the potential connectivity of the microbiota in the female urogenital system and provided insight into the exploration of urethra and genital tract diseases.

## Data Description

### Purpose of data acquisition

The role of microbiota played in the vaginal environment has received a lot of attention over the past decade, while the female upper reproductive tract was traditionally believed to be sterile and mostly studied in the context of infections or incontinence [1]. Recently, Chen et al. has determined that the female upper reproductive tract is colonised by a microbiota that can be associated with lifestyle, pregnancy history and gynaecological diseases. Like the female upper reproductive tract [2], the sterile hypothesis of urine has also been overturned by emerging evidences that indicates the existence of microorganisms in the urinary tract by cultural or sequencing approaches [3,4]. A recent study by Thomas-White *et al.* (2018), using an expanded quantitative urine culture in combination with whole-genome sequencing, has isolated and genome sequenced 149 bacterial strains from catheterized urine of both symptomatic and asymptomatic peri-menopausal women, and also showed highly similar strains of commensal bacteria in both the bladder and vagina of the same individual [5]. Gottschick *et al.* (2017) analysed the urinary microbiota of 189 individuals using 16S rRNA gene amplicon sequencing, and suggested the urethra and bladder can harbor microbial communities distinct from the vagina [6]. However, the relationship between female urine microbiota and the upper reproductive tract microbiota has so for not been studied.

Here we present a dataset of the urinary microbiota for a relatively large cohort of 147 women of reproductive age. Together with our recently published peritoneal fluid, uterine and vaginal samples from the same individuals, this data shows that although with greater amounts of *Lactobacillus* and *Streptococcus*, the urinary microbiota is more similar to the microbiota of the cervix and uterine cavity, in accordance with the anatomical opening of the bladder. Together with a wealth of collected metadata, we demonstrate this data is useful to explore the potential of the urinary microbiota for clinical diagnosis.

## Methods

### Sample collection

In this study, a total of 147 reproductive age women (age 22-48) were recruited by Peking University Shenzhen Hospital (**Supplementary Table 1**). All participants were reproductive age women who underwent hysteroscopy and/or laparoscopy for conditions without infections, such as hysteromyoma, adenomyosis, endometriosis, or salpingemphraxis. Subjects with other related diseases, such as vaginal inflammation, severe pelvic adhesion, endocrine or autoimmune disorders were removed. Pregnant women, breastfeeding women and menstrual women at sampling were also excluded. None of the subjects received any hormone treatments, antibiotics or vaginal medications within a month of sampling. In addition, no cervical treatment was performed within the previous 7 days, no vaginal douching was performed within 5 days, and no sexual activity was performed within at least 2 days.

137 self-sampling morning mid-stream urine samples were collected between December 2013 and July 2014 prior to the surgery (**Table S1**), and then stored at −80°C until transported on dry ice to BGI-Shenzhen for sequencing. The samples from an additional 10 women were collected for validation purposes by a doctor during the surgery in July 2017. For each operation, a urine catheter was inserted into the urethra to collect mid-stream urine, and same amount of sterile saline which collected through the urine catheter was set as control samples. The samples were then placed at 4°C, transported to BGI-Shenzhen, and processed within 6 hours. A portion of each sample was used for culturing live bacteria and the rest was used for sequencing.

### DNA extraction of urine samples

Genomic DNA extraction was carried out following the steps: TE buffer, lysozyme solution, proteinase K, 10% SDS were added to each urine sample, and vortexed to mix. Sample was then incubated at 55 °C in a shaking water bath for 120 min to effect complete lysis. Added phenol-chloroform-isoamylalcohol (25:24:1) to the supernatant and vortexed to mix well. Then the supernatant was transferred to a nuclease-free microfuge tube, added chloroform-isoamylalcohol (24:1) and vortexed to mix well. After the supernatant was transferred to a nuclease-free microfuge tube, sodium acetate, glycogen and cold isopropanol were added to the supernatant, and then the mixture was incubated at −20 °C for the night. 70% ethanol was used to wash the precipitate twice. TE buffer was then added to resuspend the precipitate. Extracts were treated with DNase-free RNase to eliminate RNA contamination. DNA quantity was determined using NanoDrop spectrophotometer, Qubit Fluorometer (with the Quant-iTTMdsDNA BR Assay Kit), and gel electrophoresis.[7]

### 16S rRNA amplicon sequencing

The primers 515F and 907R were utilized for PCR amplification of the hypervariable regions V4-V5 of the bacterial 16S rRNA gene. The 907R primer include a unique barcoded fusion. The primers were as follow: 515F: 5’-GTGCCAGCMGCCGCGGTAA- 3’ and 907R: 5’ - CCGTCAATTCMTTTRAGT -3’, where M denotes A or C and R denotes purine. The condition of PCR amplification was as follows: 3 min of denaturation at 94°C, followed by 25 cycles of 45 s at 94 °C (denaturing), 60 s at 50 °C (annealing), 90 s at 72 °C (elongation), with a final elongation for 10 min at 72 °C. The amplification products were purified by the AxyPrep™ Mag PCR Clean-Up Kit (Axygen, USA). The amplicon libraries were constructed with an Ion Plus Fragment Library Kit (Thermo Fisher Scientific Inc.) [8], then sequenced by Ion PGM™ Sequencer with Ion 318™ Chip v2 with a read length of 400bp (Thermo Fisher Scientific Inc., Ion PGM™ Hi◻Q™ OT2 Kit, Cat.No: A27739; Ion PGM™ Hi-Q ™ Sequencing Kit, Cat.No: A25592)[9].

### Processing of sequencing reads

The raw sequencing reads were firstly subjected to Mothur (Mothur, RRID:SCR_011947;V1.33.3) [10] for filtering of the low-quality reads utilizing the following criteria: 1) reads with less than 200bp; 2) reads not matching the degenerated PCR primers for up to two errors; 3) reads with an average quality score less than 25. A total of 8,812,607 reads, with an average of 57,225 reads per sample (a minimum of 1,113 reads and a maximum of 194,564 reads) were obtained. Subsequently, the sequences with identity greater than 97% were clustered into Operational Taxonomic Units (OTUs) using QIIME (QIIME, RRID:SCR_008249; V1.8.0) uclust programme [11], where each cluster was thought of as representing a species. The seed sequences of each OTU were aligned against the Greengene reference sequences (gg_13_8_otus) for annotation using the UCLUST taxonomy assigner.

We also calculated the Unifrac distance using QIIME based on taxonomic abundance profiles at the OTU level [11].

### PERMANOVA on the influence of phenotypes

Permutational multivariate analysis of variance (PERMANOVA) was used to assess the effect of different covariate based on the relative abundances of OTUs of the samples [12,13] using Bray-Curtis and UniFrac distance and 9999 permutations from vegan package(vegan, RRID:SCR_011950) in R [13,14].

### Real-time quantitative PCR

We quantified the four *Lactobacillus* species, including *L.iners*, *L jensenii*, *L. crispatus* and *L. gasseri* using the modified real-time quantitative PCR [15]. SYBR Premix Ex Taq GC (TAKARA) was applied to run on StepOnePlus Real-time PCR System (Life Technologies). PCR reaction mixture contained 10 μl of 2×SYBR Premix Ex Taq GC, 0.2 μM forward primer, 0.2 μM reverse primer, 1.6 μl of DNA sample and 8.2 μl ultrapure water to make up the final reaction volume of 20 μl. Each run included a standard curve and all samples were amplified in triplicate. Ultrapure water was included as blank control template.

To construct the standard curves, the sequencing-confirmed plasmids of four species were used after quantification with Qubit Fluorometer and serial 10-fold dilutions. The amplification efficiency was (100**±**10) % and linearity values were all **≥**0.99.

### Bacterial culturing

The urine samples and controls from additional 10 subjects were cultivated in the laboratory by spreading 100 μl of sample on the different agars with 5% horse blood, such as PYG agar (DSMZ 104 medium), BHI agar and EG agar. The plates were incubated in both aerobic and anaerobic conditions at 37 °C for 72 hours. To keep the medium anaerobically during culture, resazurin and cysteine-HCl were added as reducing agents. The genomic DNA of the isolates was extracted by the Bacterial DNA Kit (OMEGA), and then 16S rRNA gene amplification by the universal primers 27F/1492R were performed [16]. The amplicons were purified and sent for Sanger sequencing. The generated sequences were then submitted to BLAST on the EzBioCloud [17] for identification.

### Preliminary analysis and validation

### Microbiota composition of the urine

To explore the urinary microbiota in this dataset, morning midstream urine (UR) was self-collected prior to surgery from an exploratory study cohort of 137 Chinese women recruited (median age 31.6, range 22-48). As with our previous vagino-uterine microbiota study [2], all volunteers had conditions that were not known to involve infections (**Table S1**). From 95 women in the cohort, six locations within the female reproductive tract, including lower third of vagina (CL), posterior fornix (CU), cervical mucus (CV), endometrium (ET), left and right fallopian tubes (FLL and FRL), and peritoneal fluid (PF) were also sampled. Their vagino-uterine microbiota information has been published previously [2]. After 16S rRNA gene amplicon sequencing, the sequencing reads were pre-processed for quality control and filtering, then clustered into operational taxonomic units (OTUs) (**Methods**, **Table S2, S3**).

Due to the anatomical structures, voided urine samples from women were considered to be easily contaminated by the microbiota from the surrounding vulvovaginal region [18]. Most vaginal communities (88%) in this cohort were dominated by one genus that constitutes >50% relative abundance in the individuals. In contrast, the urinary microbiota in this study showed more heterogeneity. 56.93% of the cohort harboured a diverse type, which was represented by a significant part of the bacteria, including *Streptococcus*, *Lactobacillus*, *Pseudomonas*, *Staphylococcus*, *Acinetobacter* and *Vagococcus*, while dominated (>50%) by none of them (**Fig. 1**). In addition, 22.63% of the women harboured over 50% *Streptococcus*, and 13.87% of the women contained >50% *Lactobacillus* (**Fig. 1a, Fig. 1b**). Rare subtypes such as *Enterococcus* (2.19%), Bifidobacteriaceae (1.46%), *Prevotella* (0.73%), Enterobacteriaceae (0.73%), Coriobacteriaceae (0.73%), and *Veillonella* (0.73%) were also detected in this cohort (**Fig. 1a, Fig. 1b**). Notably, the median relative abundance of *Lactobacillus*, *Pseudomonas* and *Acinetobacter* in the urine samples were more similar to the uterus samples (**Fig. 1c**) [2]. At the phylum level, urinary microbiota was dominated by Firmicutes and Proteobacteria (**Fig. 1c**).

**Fig. 1.**
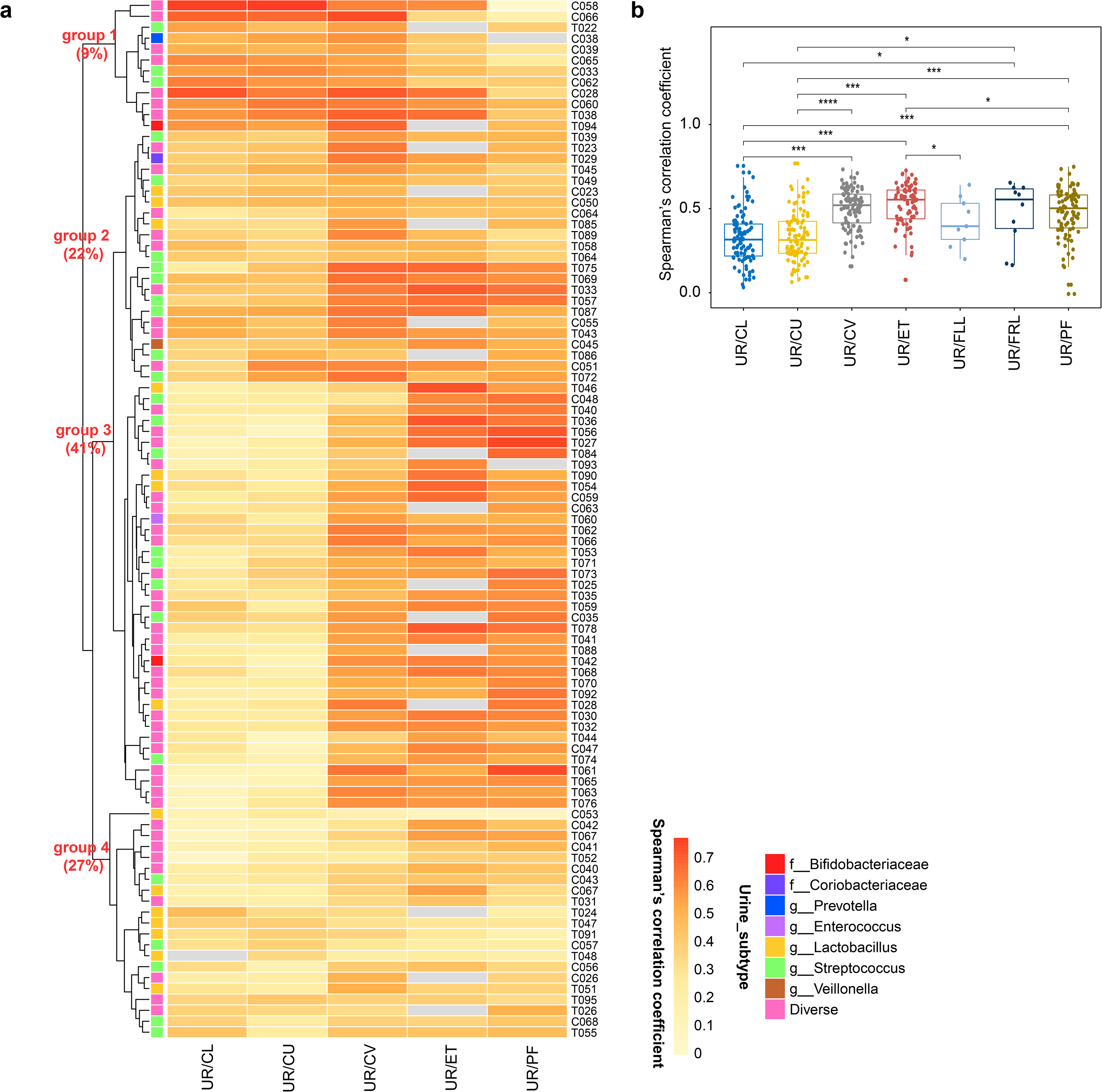
Urinary microbiota of the initial cohort of 137 Chinese reproductive-age women. **(a)** The relative abundances of genera detected in each individual are shown in the bar chart. The dendrogram is a result of a centroid linage hierarchical clustering based on Euclidean distances between the microbial composition proportion of urinary bacterial communities. (b) The ratio of different urinary microbiota types. The genus whose relative abundance account for >50% in an individual is selected as an identified type. The genera all account <50% of the microbiota in an individual are identified as diverse type. (c) Pie chart for the urinary microbial genera according to the median relative abundance. Genera that took up less than 1% of the microbiota were labelled together as ‘others’. The outer-ring indicates the distribution of microbiota in phylum level (**Table S1**).

### Intra-individual similarity in the urine-reproductive tract microbiota

To further assess the microbiota relationship between the urine and the six positions of the reproductive tract, we computed the intra-individual correlation of microbial profile between the urine and different sites of the reproductive tract and clustered the individuals into 4 groups (Spearman correlation coefficient, **Fig. 2a, Table S4**). Interestingly, the microbiota of group 3, which accounted for 41% of the cohort, showed the significance of correlation with urine samples increased from CL to CV, ET and PF gradually. In contrast, 9% women in group 1 presented a reverse trend. In group 2 (22%) and group 4 (27%), there appeared to be a weak relationship between the urine and reproductive tract. Taken together, we observed the most similar distribution of microbiota between urine and CV/ET (**Fig. 2b**). The principal coordinate analyses (PCoA) of the weighted and unweighted intra-individual UniFrac distance further corroborated our results that there is an intra-individual similarity of the microbiota between the urine and the upper sites of reproductive tract, especially the junction sites (CV and ET) (**Fig. 3**).

**Fig. 2.**
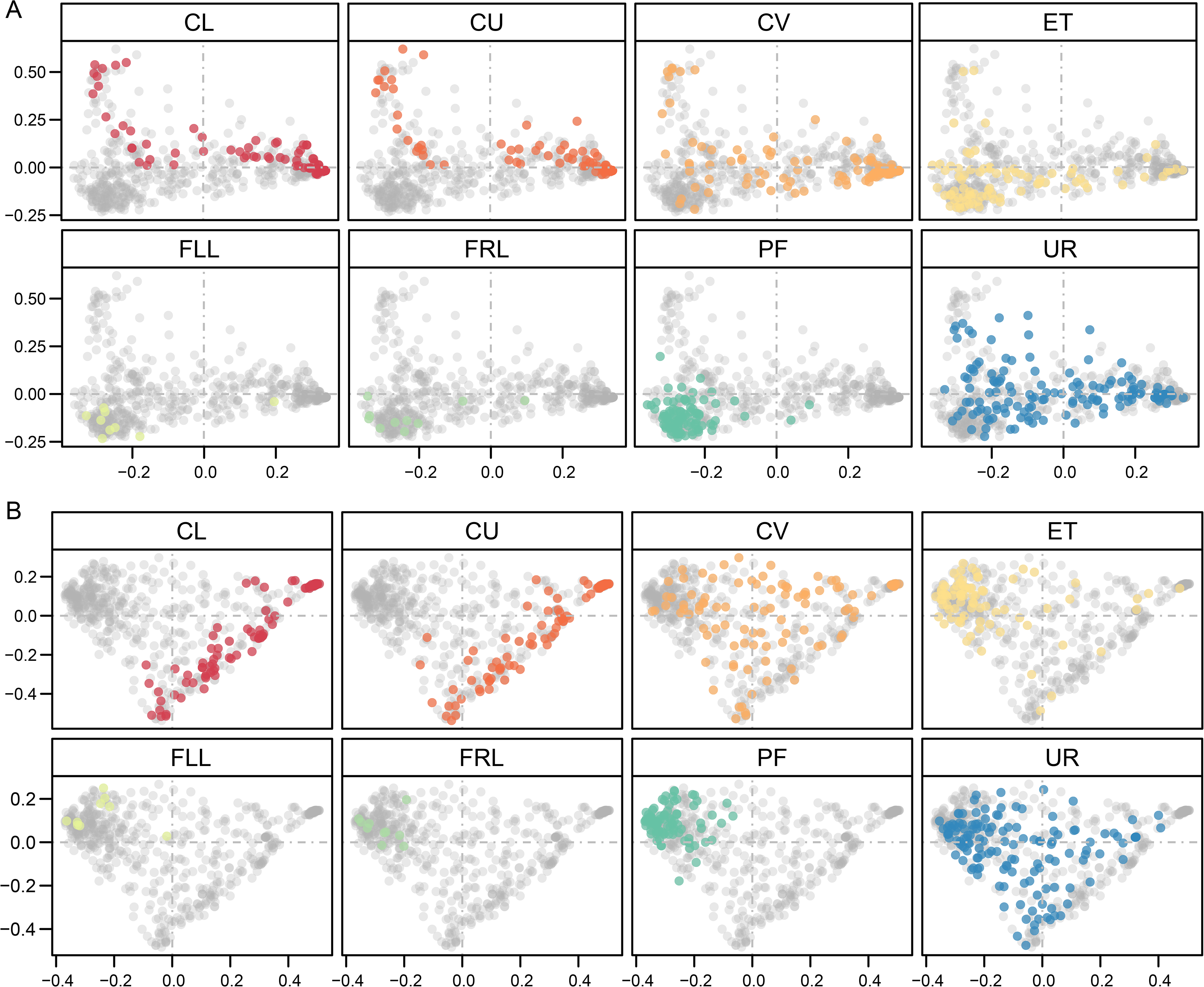
Similarity of the urine-reproductive tract microbiota within individuals. (**a**) Heatmap for the intra-individual Spearman’s correlation coefficient between urine and different body sites of reproductive tract (Table S4). Samples derive from the initial cohort of 95 Chinese reproductive-age women, who collected both the urine and reproductive tract samples. As the number of samples from fallopian tubes (FLL, FRL) is too small, the correlations between urine and fallopian tubes are not shown. The dendrogram is a result of a centroid linage hierarchical clustering based on Euclidean distances between the intra-individual Spearman’s correlation coefficient of different body sites. The colored squares illustrate the subtypes of urinary microbiome. (**b**) Spearman’s correlation coefficient between urine and different body sites of reproductive tract. Wilcoxon ranked sum test is used to calculate the difference. Boxes denote the interquartile range (IQR) between the first and third quartiles (25th and 75th percentiles, respectively), and the line inside the boxes denote the median. The whiskers denote the lowest and highest values within 1.5 times the IQR from the first and third quartiles, respectively. An asterisk denotes p <0.05, two asterisks denote p <0.01, three asterisks denote p <0.001.

**Figure 3.**
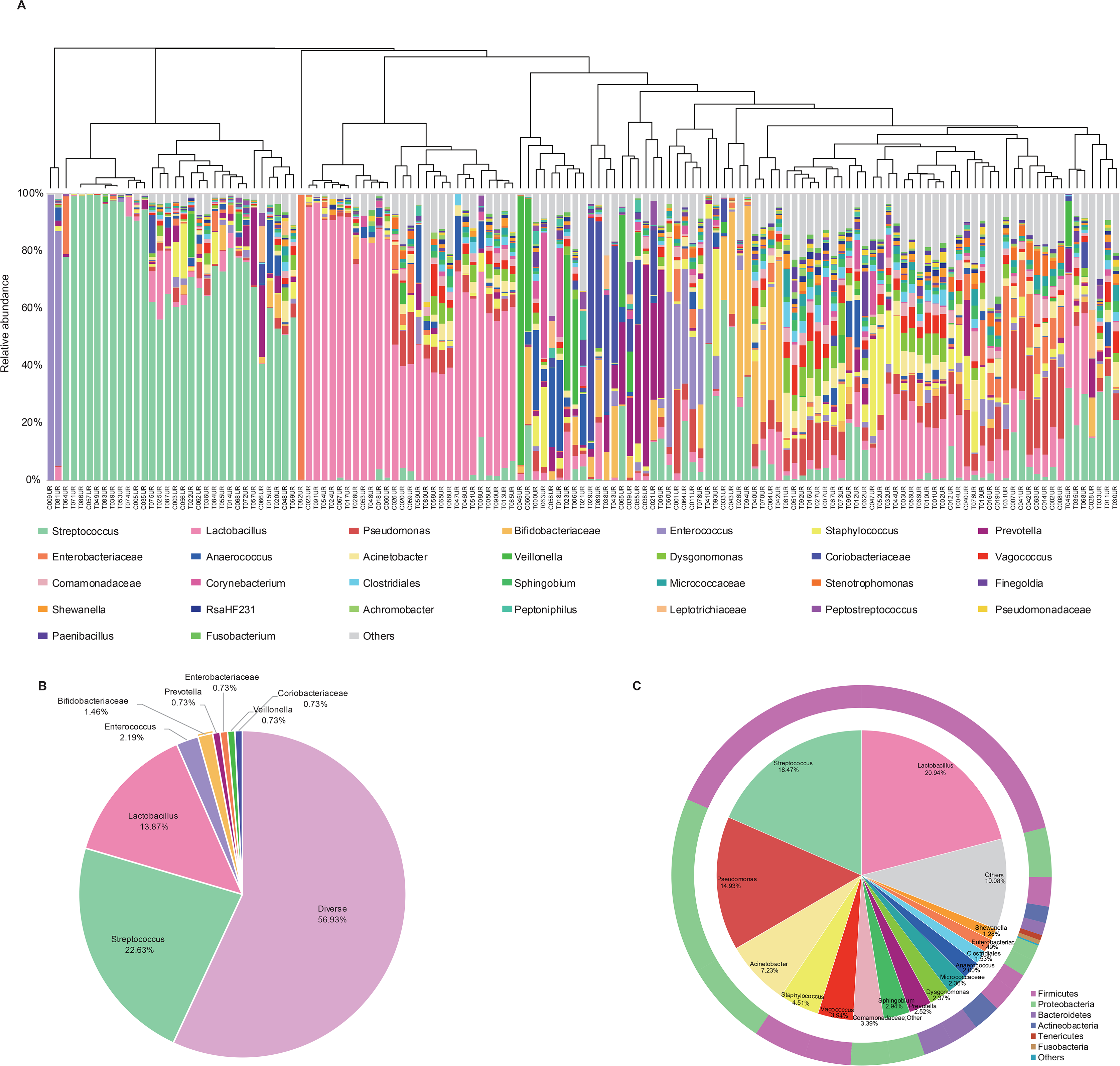
PCoA on the samples based on Unweighted-UniFrac (a) and Weighted-UniFrac (b) distances. Samples were taken from UR, CL, CU and CV before operation, and from ET and PF during operation. Samples derive from the initial cohort of 137 Chinese reproductive-age women. Each dot represents one sample (nLJ=LJ94 CL, 95 CU, 95 CV, 80 ET, 93 PF, 9 FLL, 10 FRL and 137 UR).

### Lifestyle and clinical factors influencing the urinary microbiota

The human microbiome is dynamic and highly affected by its host environment. Age, menstrual cycle, benign conditions such as adenomyosis, and infertility due to endometriosis have previously been reported to shape the vagino-uterine microbiota [2]. With our comprehensive collection of demographic and baseline clinical characteristics from women of reproductive age (**Table S1**), such variations in the urinary microbiota can be explored in this dataset. Urinary microbial composition was significantly associated with these factors, such as age, surgical history, abortion, vaginal deliveries, experience of given birth (multipara vs. nullipara), infertility due to endometriosis and hysteromyoma (PERMANOVA, P < 0.05, q > 0.05, **Table 1**). Although the urinary microbiota also correlated with some other factors, such as menstrual phase, contraception, endometriosis, pelvic adhesiolysis and anemia, the statistical significance was not achieved after controlling for multiple testing (PERMANOVA, P < 0.05 but q > 0.05, **Table 1**). The initial results here indicate a close link between the urinary microbiota with the general and diseased physiological conditions, and can be taken further with further exploration of this data.

### Potential uses

As the large-scale cohort for studying the female urinary microbiota, our data provide a useful baseline and reference dataset in women of reproductive age. We also explored the association between the composition of urinary microbiota and that of reproductive tract microbiota. It is valuable to note that a higher intra-individual compositional similarity of the microbiota between urine and cervical canal/uterus instead of the vagina, which indicate that sampling of midstream urine (the least invasive and the easiest way) could be potentially used to survey the micro-environment of the cervical canal and uterus in the general population. This is of relevance in relation to the demonstrated associations between the urinary microbiota and various uterine-related diseases such as hysteromyoma, and infertility due to endometriosis. Our data provide a reference for clinical diagnosis, and warrants further detailed exploration. Thus, we hope this dataset will be the beginning of a new round of accelerated discoveries, including a novel scientific explanation of the uterine-related diseases by longitudinal studies on the microbiota of urinary and reproductive tract.

## Declarations

### Ethics approval and consent to participate

The study was approved by the Institutional Review Board of BGI-Shenzhen and Peking University Shenzhen Hospital. All participants gave written informed consent prior to their recruitment into the study.

### Availability of supporting data

The sequence reads generated by 16S rRNA gene amplicon sequencing have been deposited in both the European Nucleotide Archive with the accession number PRJEB29341 and the CNSA (https://db.cngb.org/cnsa/) of CNGB database with accession code CNP0000166. The sequences of bacterial isolates have been deposited in the European Nucleotide Archive with the accession number PRJEB36743

## Supporting information

Additional finding from sampling of this dataset and the related figure legends

## Acknowledgments

The study was supported by the Shenzhen Municipal Government of China (JCYJ20160229172757249 and JCYJ20170817145523036) and Shenzhen Peacock Plan (No. KQTD20150330171505310). The authors really appreciate colleagues at BGI-Shenzhen for DNA extraction, library construction, and sequencing.

## Author contributions

H.J. and R.W. organized this study. W.W., J.D., H.D., L.Z., H.T., T.W and R.W. performed the sample collection, and phenotypic information collection. F.L., L.S., C.C. and J.L. performed the molecular biology experiments. C.C., L.H. and F.L. performed the bioinformatic analyses. C.C., X.Z., F.L. and H.J., wrote the manuscript.

