## Additional finding from sampling of this dataset and the related figure legends for "The female urinary microbiota in relation to the reproductive tract microbiota"

Cultivation of live bacteria from Transurethral catheterized urine

The question of whether bacterial DNA signals are originated from live bacteria or fragments in the urine samples has aroused heated discussion. To demonstrate the utility of data for this purpose we performed a validation study using live bacteria culture of urine samples from an additional cohort of 10 women.

We tried to culture and isolate bacterial colonies from freshly collected urine samples. Urine samples were serial diluted and spread on three different kinds of agar plates and incubated under both aerobic and anaerobic conditions. Six different positive isolates belonging to 5 genera, including *Lactobacillus, Staphylococcus, Clostridium, Enterococcus and Propionibacterium* were obtained from 3 out of 10 subjects (**Table S5**). The 5 genera were also found as dominant in our 16S rRNA gene amplicon sequencing data, and consistent with previous cultivation results of published papers. [1–4] (**Table S5**). Reassuringly, no isolates were detected from the negative controls (sterile saline and ultrapure water). Therefore, this verified the existence of live bacteria in the urine by obtaining isolates using conventional culturing methods.

Considerable bacterial biomass revealed by qPCR

To provide some additional evidences of the bacterial communities in the urine, a species-specific real-time qPCR method was utilized to focus on the four common vaginal *Lactobacillus* species, i.e. *L. crispatus*, *L. iners*, *L. jensenii* and *L. gasseri* (**Fig. S1**). The *Lactobacillus* species we examined presented a similar distribution and abundance along the reproductive tract, and the corresponding urinary *Lactobacillus* ranged between upper and lower reproductive tract (**Fig. S2a**). Among them, *L. iners* occurred most frequently (59%) in the urine samples, while *L. crispatus* only occurred in 26% of women sampled (**Fig. S2b**). *L. iners* was reported far less protective against bacterial and viral infections compared to *L. crispatus* [5]. 80% of the cohort was detected to harbour at least one of these four *Lactobacillus* species (**Fig. S2b**). The occurrence rate of *Lactobacillus* in genus level of 16S rRNA gene amplicon sequencing data was 94% (**Fig. 1a**). The total bacterial biomass is approximated by the ratio of the copy number from the result of qPCR to the relative abundance according to the result of 16S rRNA gene sequencing of the same sample. The result gave an estimation of 10^7^, placing the urinary bacterial biomass between the vaginal-cervical sites (10^10-^10^11^) and the endometrium (ET) samples (10^6-^10^7^) [6] (**Fig. S2**), all of which were orders of magnitude above potential background noise [7]. These results were interestingly consistent with a weakly acidic pH of the urine, in comparison to pH < 4.5 in the vagina or pH ~ 8 in the peritoneal fluid [8].

**Supplementary figure legends**

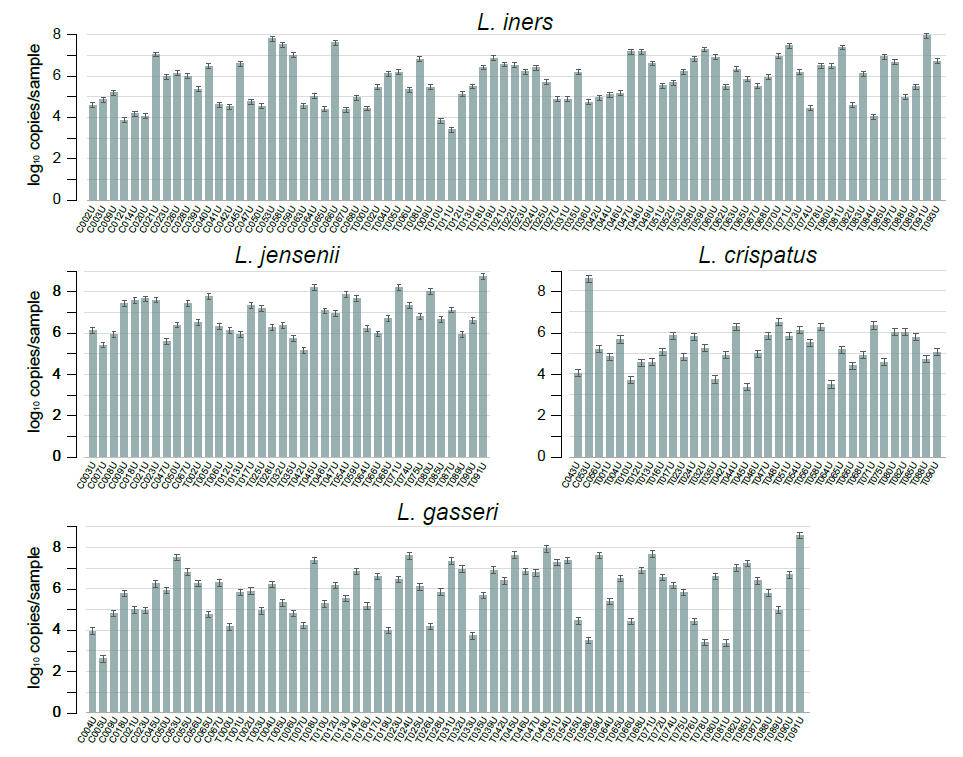

**Fig. S1. The presence and average concentrations of the dominant Lactobacillus species at urine samples determined by real-time qPCR.** (a) *Lactobacillus iners*. (b) *Lactobacillus jensenii*. (c) *Lactobacillus crispatus*. (d) *Lactobacillus gasseri*. Error bar represents the standard deviation of three replicate measurements.

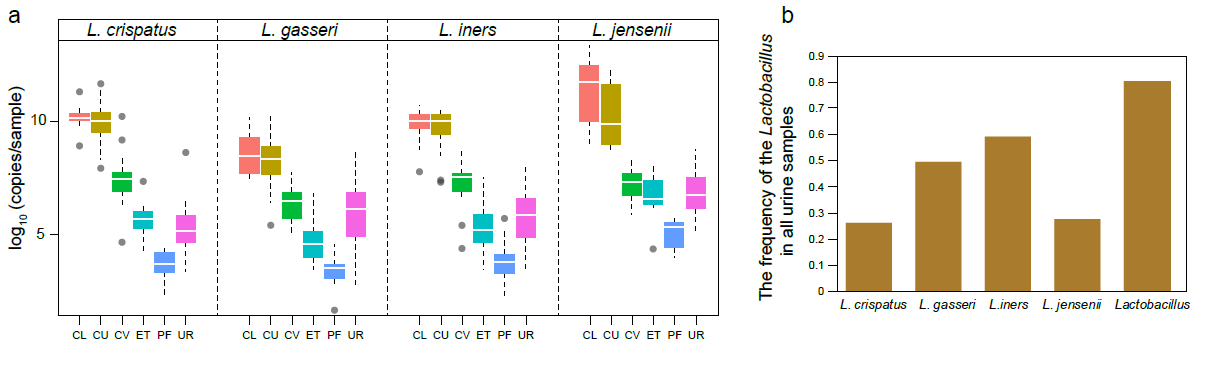

**Fig. S2. The concentrations of the dominant *Lactobacillus* species at urine and the reproductive tract.** Samples derive from the initial cohort of 137 Chinese reproductive-age women (**Table S1**). (**a**) The abundance of *Lactobacillus iners*, *Lactobacillus jensenii*, *Lactobacillus crispatus* and *Lactobacillus gasseri* calculated by qPCR results in different samples. Boxes denote the interquartile range (IQR) between the first and third quartiles (25th and 75th percentiles, respectively), and the lines inside the boxes denote the median. The whiskers denote the lowest and highest values within 1.5 times the IQR from the first and third quartiles, respectively. (**b**) The frequency of the respective *Lactobacillus* detected in all urine sample.
